# Bioconversion of Mango Pulp Industrial Waste into Ellagic acid Using *Aspergillus niger*

**DOI:** 10.1101/2020.03.17.996074

**Authors:** Athiappan Murugan, Anandan Rubavathi, Visali Kannan, Aurumugam Parthiban

**Affiliations:** Department of Microbiology, Periyar University, Salem 636011, India

**Keywords:** Ellagic acid, *Aspergillus niger*, Bioconversion, mango pulp waste, solid-state fermentation

## Abstract

Ellagic acid was considered as the potential bioactive compound with many therapeutical applications. Bioconversion of tannin present in the mango pulp processing waste in to ellagic acid using fungi would be better alternate than the chemical as well as extraction from plant sources. A total of three different fungi were isolated from the soil sample and it was confirmed as *Aspergillus niger*. Further, the isolated strains of *A. niger* were identified to produce ellagic acid from ellagitannin of mango waste. Quantification of the ellagic acid production was carried out by solid-state fermentation using 3% of mango waste as substrate. Ellagic acid enzyme activity was calculated and found to be 17.6 U ml^−1^ The ellagic acid production was optimized to fix the various factors, that is, pH and temperature, nitrogen and carbon source. The maximum production (200 μg/g) of ellagic acid was achieved at pH 5.5, temperature 30 °C, Ammonium nitrate as nitrogen source, 0.2% of NaCl and carbon source (0.2% of sugar) with 3% of mango pulp waste. Ellagic acid produced was characterized by UV–vis spectrophotometer and by FT-IR analysis.

## Introduction

Natural products are used widely to cure many human diseases. Ellagic acid (EA) is a phenolic compound with potent anti-oxidant, anti-carcinogenic, anti-viral, anti-mutagenic, anti-parasitic, anti-diabetics, anti-steatosic, anti-cholestic, anti-fibrogenic and anti-hepatocarcinogenic properties [1,8]. However, ellagitannins inhibit the growth of a number of bacterial species and resist microbial attack. But certain moulds, such as *Aspergillus* sp., *Penicillium* sp., *Trichoderma* sp., *Fusarium* sp., *Mucor* sp., *Rhizopus* sp. *, Neurospora* sp., are able to hydrolyse tannin with the help of tannases [9]. Hydrolyse volania tannin to ellagic acid have been demonstrated with *Aspergillus niger* grown on waste husk [10].

The efficiency of EA as antioxidant compounds greatly depends on their chemical structure and several hydroxyl groups of ellagic acid found to enhance their activity [11]. The anti-oxidant efficiency of EA is also depending on their degree of hydroxylation. Ellagic acid is a dilactone formed by the hydrolysable of hexahydroxydiphenic acid. EA is found in several plants, such as oak tree, eucalyptus, pomegranate, strawberry, raspberry, blueberry, blackberry, cranberry, gooseberry, grape, pecan, walnut, valonea and creosote bush at varying concentration [12]. These ellagic acids are being extracted from ellagitannin-rich plant sources using strong alkali and acids (H_2_SO_4_ or HCl) [13, 14]. It also involves treatment with higher concentrations of alkali or acid that leads to corrosion or damage of the process vessels and subsequently needs more safety precaution. Major disadvantage is the unforeseen toxic or hazardous with chemically produced [15].

Biotransformation of ellagitannin to ellagic acid is known for decades and found economic. Biosynthesis of ellagic acid achieved by the enzymatic catalysis of ellagitannin by the tannase. Tannase acts on depside (C–O) and ester linkages present in hydrolysable tannins resulting into a biotransformed products such as gallic acid, ellagic acid and glucose [16]. Microbial synthesis of ellagic acid using solid-state fermentation with mango waste by the *Aspergillus* sp., a GRAS organism (Generally Recognized as Safe) is planned in this study. Huge volume of waste generated in the mango pulp processing industries is harnessed for the production of ellagic acid. Depending on the varieties, the kernel represents 45–85% of the seed containing 17–22% of tannin being recycled for the production of valuable ellagic acid [17]. Optimization of ellagic acid production through onsite fermentation would be great boon to the smaller industries.

## Materials and methods

### Isolation and identification of *A. niger*

Soil samples were collected from Periyar University campus, Salem, Tamil Nadu. The soil samples were collected in sterile sealed polythene bags using sterile spatula. One gram of soil sample was dispensed into 100 ml of sterile distilled water and shaken well for 20 min and serially diluted was carried out up to 10^−7^ ml. One ml of each serially diluted sample was poured in the sterile petri plates containing Potato Dextrose Agar (PDA) medium. Chloramphenicol (25 μg/ml) was added in molten PDA medium in the petri plates and was incubated at 30 ± 1°C for 3–5 days for the isolation of *A. niger*. After incubation, different fungal colonies grow on the PDA medium. Preliminary identification of the fungi was done based on their morphological characteristics. Microscopic examination of fungi done by placing one drop of Lactophenol Cotton Blue (LPCB) on glass slide and immersed the fungal culture in a drop and placed cover slip on the slide. The slide was observed under light microscope (40× objectives).

### Sub-culturing of *A. niger*

Purified culture of *A. niger* strain was sub-cultured in Potato Dextrose Agar (PDA) plates. Once a week, the culture was sub-cultured in fresh medium.

### Screening for Ellagic acid producer

Czapek minimal media was prepared and sterilized and the solution of ellagitannin was sterilized separately by passing through a membrane filter (pore size 0.22 μm, Millipore Corporation, Bedford, MA, USA) and was added to Czapek Dox’s minimal medium to a final concentration of 1%. Fungi were inoculated on czapek medium containing ellagitannin as sole carbon. The plates were incubated at room temperature at 4–5 days. After the incubation zone formation around the fungal growth was observed. If zone formation was not clear the plate flooded with 0.5M FeCl_2_ which reacted with the tannin-protein complex, yielding a brown to black color, providing a dark background allowing clear visibility of zones of ellagitannin degradation.

### Preparation of fungal spore suspension

Spore suspension was prepared a by pouring 5–7 ml of sterile 0.9 % NaCl (w/v) + 0.01 % Tween 80 on a PDA plate containing *A. niger* culture as in sub-heading and scrape the spores into solution. Filter the spore solution into a sterile 10 ml test tube through a sterile funnel containing a cotton wool plug to remove hyphae. The number of spores can be counted using haemocytometre and diluted to 2 × 10^7^ spores/ml.

### Production of Ellagic acid

About 3 g of mango pulp wastes was taken in 250 ml conical flask and 7 ml of Czapek Dox medium in test tubes were autoclaved separately at 121 °C for 15 min at 15 lbs. After sterilization, 7 ml medium was poured into conical flask containing mango pulp wastes at 3%. One millilitre of inoculum (containing 2 × 10^7^ spores/g of support) was inoculated, and flasks were incubated at 30 °C and were monitored kinetically for 144 h.

### Extraction of ellagic acid

After fermentation, the biomass was mixed with distilled water at (1:2) ratio heated for 5 min. After cooling, it was centrifuged at 10,000 rpm and the supernatant was measured. Diethyl ether (5×) was added to the supernatant and mixed thoroughly by swirling for 2 min. The solution was allowed to stand for 2 min at room temperature for the separation of the organic phase and aqueous phase. In the biphasic system, ellagic acid was extracted into the organic phase, which was then separated and the residue was used for further analysis.

### Quantification UV–vis spectrophotometer

Stock solution of standard ellagic acid (50 mg/50 ml) was prepared by dissolving in methanol. Working standards were prepared by tenfold dilution from this working standard pipette out 0.2 ml, 0.4 ml, 0.6 ml, 0.8 ml, 1.0 ml, 1.2 ml, 1.4 ml, 1.6 ml, 1.8 ml and 2.0 ml into the clean test tubes and the tubes was appropriately labeled. The tubes were making up to 2 ml using phosphate buffer (0.05 M pH 7.4), and optical density (OD) was read at 255 nm. A standard graph was prepared and floating concentration of standard ellagic acid with OD values. The amount of ellagic acid produced in the biomass was determined using standard graph.

### Determination of tannase activity

One millilitre of crude enzyme liquid was added to 1 ml of 23% (v/v) solution of gallotannin which replaced the wheat bran as substrate in a citrate buffer (0.1 M pH 5.0). After incubation at 40 + 1 °C for 10 min, the reactions were stopped by the addition of 1 ml ethanol (90%), and absorbance was measured at 270 nm. Control samples were produced by adding 1 ml of crude enzyme to the same reaction mixture already containing the ethanol, and these samples were withdrawn after incubation at 40 + 1 °C for 3 min. The quantity of gallic acid released during hydrolysis of gallotannin represents the tannase activity. Here, one unit of tannase activity corresponds to a decrease of 0.001 units in the absorbance per milliliter of gallotannin solution per minute at 270 nm.

### Characterization of ellagic acid

#### FT-IR analysis

Fermented broth containing ellagic acid was transferred to centrifuge tubes, and ellagic acid was separated by centrifugation at 2000 rpm for 20 minutes. The supernatant was transferred to separating funnel, and the equal volume of ethyl acetate was added to the supernatant and mixed well for separation. The upper aqueous layer containing ellagic acid was collected in a beaker. Then it was allowed for evaporation of ethyl acetate. About 2 ml of methanol was added after evaporation and mixed well and followed for storage in refrigerator for further experimental analysis using Fourier transform infrared spectroscopy (FT-IR).

### Optimization of ellagic acid production

#### Effect of temperature

The solid-state fermentation was carried out at different temperatures of 25 °C, 30 °C, 35 °C and 40 °C for 144 h.

#### Effect of pH

The solid-state fermentation was carried out at different pH (adjusted using 1 N NaOH and 1 N HCl) ranging from 5.0, 5.5, 6.0 and 6.5.

#### Effect of nitrogen supplements

The effect of nitrogen sources was studied on production of ellagic acid. About 0.2% of various nitrogen sources such as sodium nitrate, sodium nitrite, ammonium nitrate, ammonium chloride, ammonium sulfate, potassium nitrate and urea were added to fermentation media.

#### Effect of mango pulp supplements as carbon source

Acid hydrolyzed mango pulp waste containing 45–65% of sugar was used as a carbon source [18].

#### Mango pulp waste pretreatment

Acid hydrolysis of mango pulp waste performed by 50 ml of 1% hydrochloric acid heated with 5 g of mango pulp at 120 °C at 10 min. After heat treatment, the suspension was centrifuged at 5000 rpm for 10 min, and the supernatant was collected and analyses the amount of glucose content by DNS method.

#### Reducing sugar estimation in mango pulp pretreated suspension

Standard solution was prepared by dissolve 100 mg of glucose in 50 ml of distilled water. From the standard solution pipette out 0.2 ml, 0.4 ml, 0.6 ml, 0.8 ml, 1.0 ml, 1.2 ml, 1.4 ml, 1.6 ml, 1.8 ml and 2.0 ml into the clean test tubes and the tubes was appropriately labeled. About 0.1 ml of mango pulp pretreated sample added into the test tubes. The tubes containing solutions were making up to 2 ml using distilled water and 2 ml of distilled water pipette out into the test tube consider as a blank. From each tube, 2 ml of DNS reagent was added and the tubes were kept in water bath for 5 min and the tubes were cooled in running tap water. The absorbance read at 520 nm, and the standard graph was plotted to detect the glucose level in sample.

#### Mass production of ellagic acid

Owing to produce large number of fungal spores, 7 ml of Czapek Dox medium in test tubes was separately autoclaved at 121 °C for 15 min at 15 lbs and supplemented with 3 g of mango waste in 250 l conical flask. One millilitre of inoculum (containing 2 × 10^−7^ spores/g of support) was inoculated, and flasks were incubated optimized temperature. Samples were monitored daily up to 6 days.

## Results and discussions

### Isolation and identification of *A. niger*

A total of three different fungal strains were isolated from the soil sample using PDA plates after 4 days of incubation. Yellow to white mycelia with black color sporulating fungus (Fig 1) was examined by using LPCB mount. Based on the microscopic (hyaline, septate hyphae, conidiophore’s were long and globose) and macroscopic morphology (presence of black, globose conidia with very dark to black spores) (Fig 2), the fungal strain has been as confirmed as *A. niger*. *A. niger* was included Generally Recognized as Safe (GRAS) under 21 CFR act of U.S. Food and Drug administration (1958), and it has been used for the ellagic acid production from Pomegranate Husk [19,20]. Conversely, group have also produced ellagic acid using solid-state fermentation from cranberry pomace with *Rhizopus oligosporus* [21].

**Fig 1:**
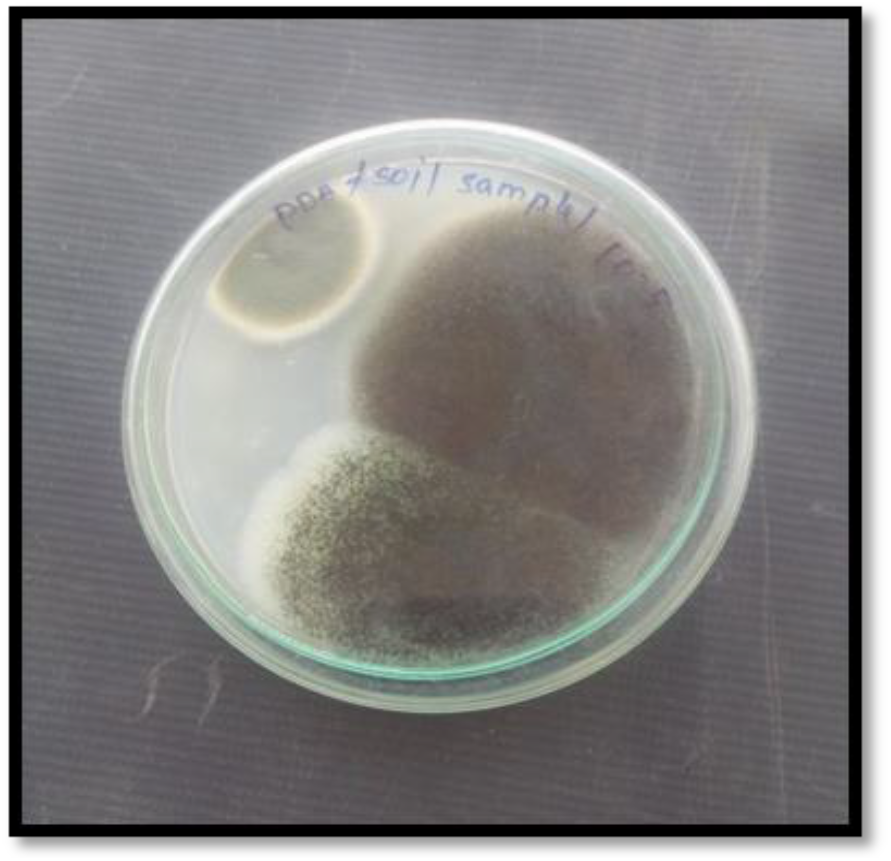
Sporulation of *A. niger* on PDA plate

**Fig 2:**
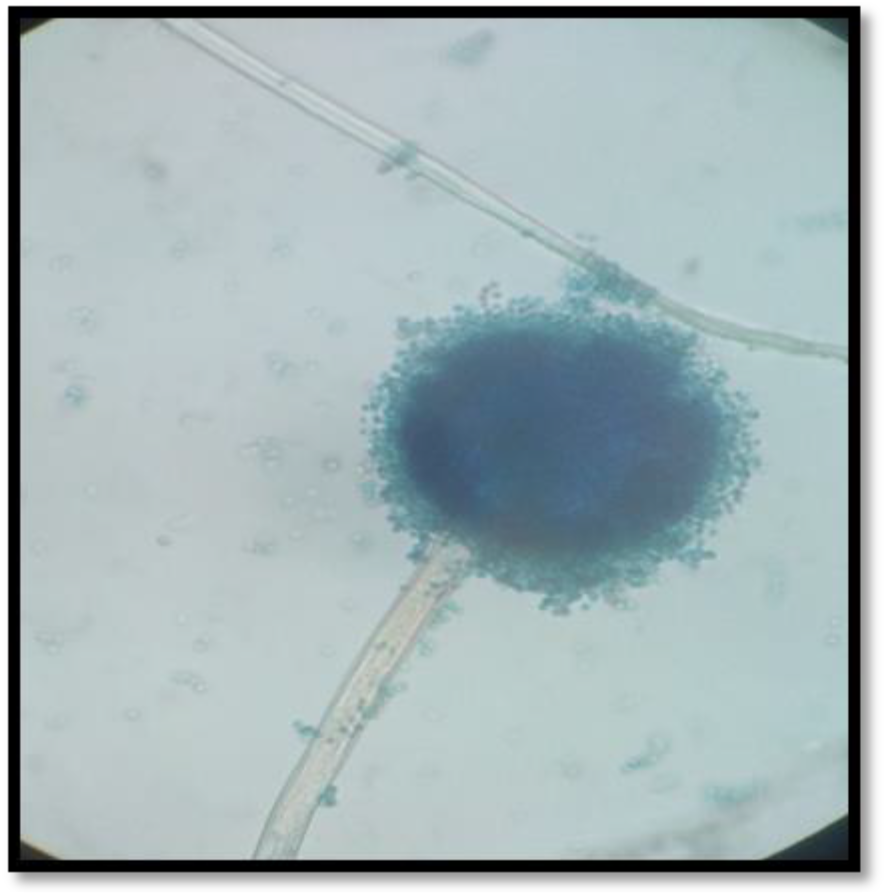
Microscopic observation of *A. niger*

### Screening of ellagic acid producer

Ellagic acid producer was identified by clear hollow zone around the fungal colonies which indicated the breakdown of ellagitannin supplemented in czapek dox minimal media. Then plates were flooded with ferric chloride which reacted with tannin to produce dark brown color and the zone was clearly visible (Fig 3). Ellagitannin acyl hydrolase was an enzyme responsible of the EA production [22]. Mycelium of potential tannase producers was scanty but clear zones were produced [23]. *Aspergillus* sp. been recorded as best extracellular tannase producing fungi [24]. Ellagic acid and gallic acid production from ellagitannin and gallotannin by *A. niger* had been proved [19,25–29].

**Fig 3:**
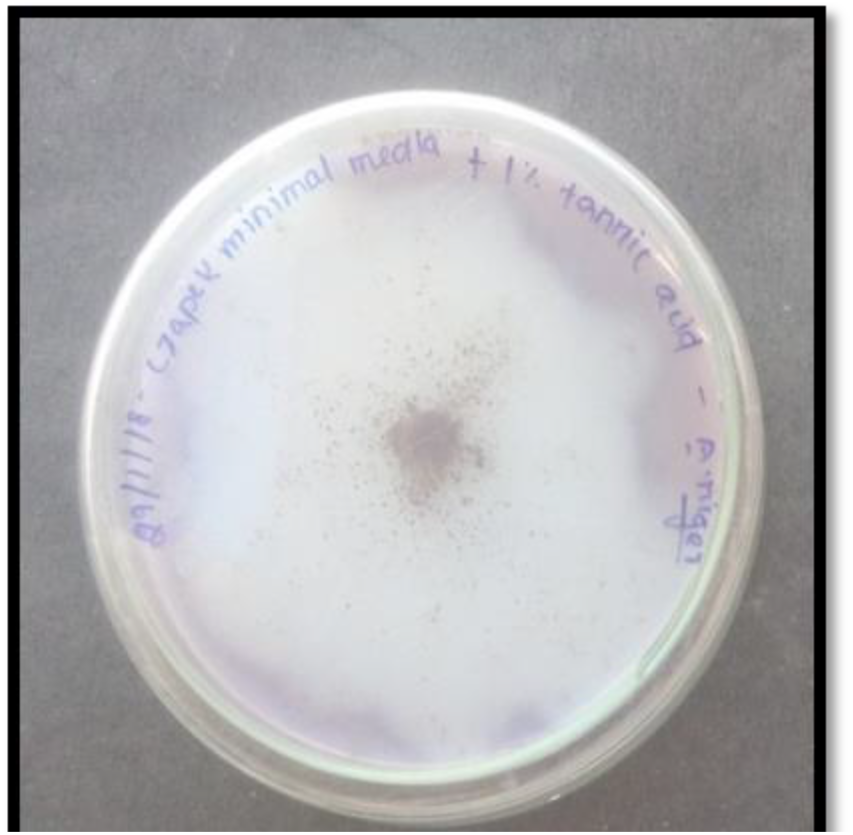
Plate showing Clear zone around ellagic acid producer on Czapek Dox Minimal Media

### Spore count and inoculum preparation

Fungal spores were dissolved in sterile solution of 0.01% Tween-80 and counted in a Neubauer chamber as prescribed [30]. Total number of spores counted in five squares was 1, 38,40,0 and the average number of spores were 27,680.

**Number of spores/ml** = Average no of spores × dilution factor × 10^4^

**Number of spores/ml** = 27680 × 1 × 10^4^

**Number of spores/ml** = 27.68 × 10^7^

One milliliter of spore suspension contains 27.68 × 10^7^ spores which were taken as stock culture. From the stock, 1 ml was pipette out and made up to 4.6 ml, to obtain 6 × 10^7^ spores/ml and used as inoculum.

### Production and quantification of ellagic acid

Ellagic acid production by *A. niger* was carried out by solid-state fermentation using mango waste as substrate. Similarly, pomegranate husk was used as a substrate for the EA production by *A. niger* [30]. Ellagitannin acyl hydrolase enzyme activity was calculated and found to be 17.6 U ml^−1^. The extracted sample containing ellagic acid was analyzed using UV–vis spectrophotometer at 255 nm and compared with standard ellagic acid. It was found that 200 μg/ml of EA produced from 3% of mango waste by *A. niger* (Chart 1).

A prior study revealed that optimum conditions for tannase production were solid to liquid ratio of 1:2, 35 °C, pH 5.5 and 72 h incubation time which resulted in 0.256 mg/ml of ellagic acid [31]. Already recorded the solid-state fermentation of pomegranate peel powder has also produced 8.48–132.62 mg/g of ellagic acid under optimized condition by *A. niger* GH1 [26]. SSF of pomegranate seeds and husk by *A. niger* GH1 and PSH to yields 6.3 and 4.6 mg of EA per gram of dried pomegranate husk [19]. However, maximum EA (138.44 mg g^−1^) production was recorded at 8 h, using *A. niger* GH1 through submerged culture [20]. Degradation of creosote bush ellagitannin yielded 23.1 % of ellagic acid. Cups extract of valonia acorns degrade by mixed culture (*Aspergillus oryzae* and *Trichoderma reesei*) to yields 23% of EA [33]. Biosynthesis of volania tannin hydrolase by *Aspergillus* SHL6 has yielded 5.0 g l^−1^ of ellagic acid [29]. *A. niger* coculture with *C. utilis* degrades volanea tannin to yield 12.1%. Ellagic acid production from mango pulp wastes was lower than the previous report because tannin content of mango pulp wastes was much lower than the used substrate.

### Characterization of extracted ellagic acid

#### FT-IR analysis

Characterization of the ellagic acid was done by using FTIR by their functional groups. Test compound was analyzed by FT-IR, the results of the spectrogram represent the presence of glycosidic groups V (O-H) that is the presence of OH stretching with a broad peak at 3554 cm^−1^. The carboxylic substituent’s C=O stretching peak at 1691.57 cm^−1^. The amide group at 1193–1057 cm^−1^ and the alkaloids at 638.44 cm^−1^ correspond to ellagic acid. All these bands are more or less similar to the standard ellagic acid. Based on the functional group and the chemical structure of the extracted compound, it may be presuming as ellagic acid. The sample was further confirmed with the comparison with standard ellagic acid (Chart 2). In cultural heritage field, FTIR studies of tanning materials have been performed in bookbinding leathers and parchment samples. These bands can be assigned to the ethereal C–O asymmetric stretching vibration arising from the pyran-derived ring structure of this class of tannins [34].

### Optimization of various factors for ellagic acid production

#### Effect of pH

Fungal growth and enzymes activity were highly sensitive to pH. Among various pH (3.0–8.0) provided for the production of EA maximal fungal growth and maximum tannase activity (18.4 U ml^−1^) was observed at pH 5.5 (Chart 3) and (Table 1). Similarly, reported that, *Aspergillus flavus* needed an acidic environment for tannase production [35]. Already suggested the tannase production from *A. niger* showed optimum activity at pH of 6.0 with a second peak at pH 4.5 [36]. Maximum tannase production in *Aspergillus* sp. was observed at pH at 5.5 [37]. Previous study reported that strains of *A. niger* possess tannin protein complex degrading activity at a pH range of 6.0 and 5.0, respectively [38]. However, reported the optimum pH for tannase production from *R. oryzae* was at pH 5.0 [39].

**Table 1.**
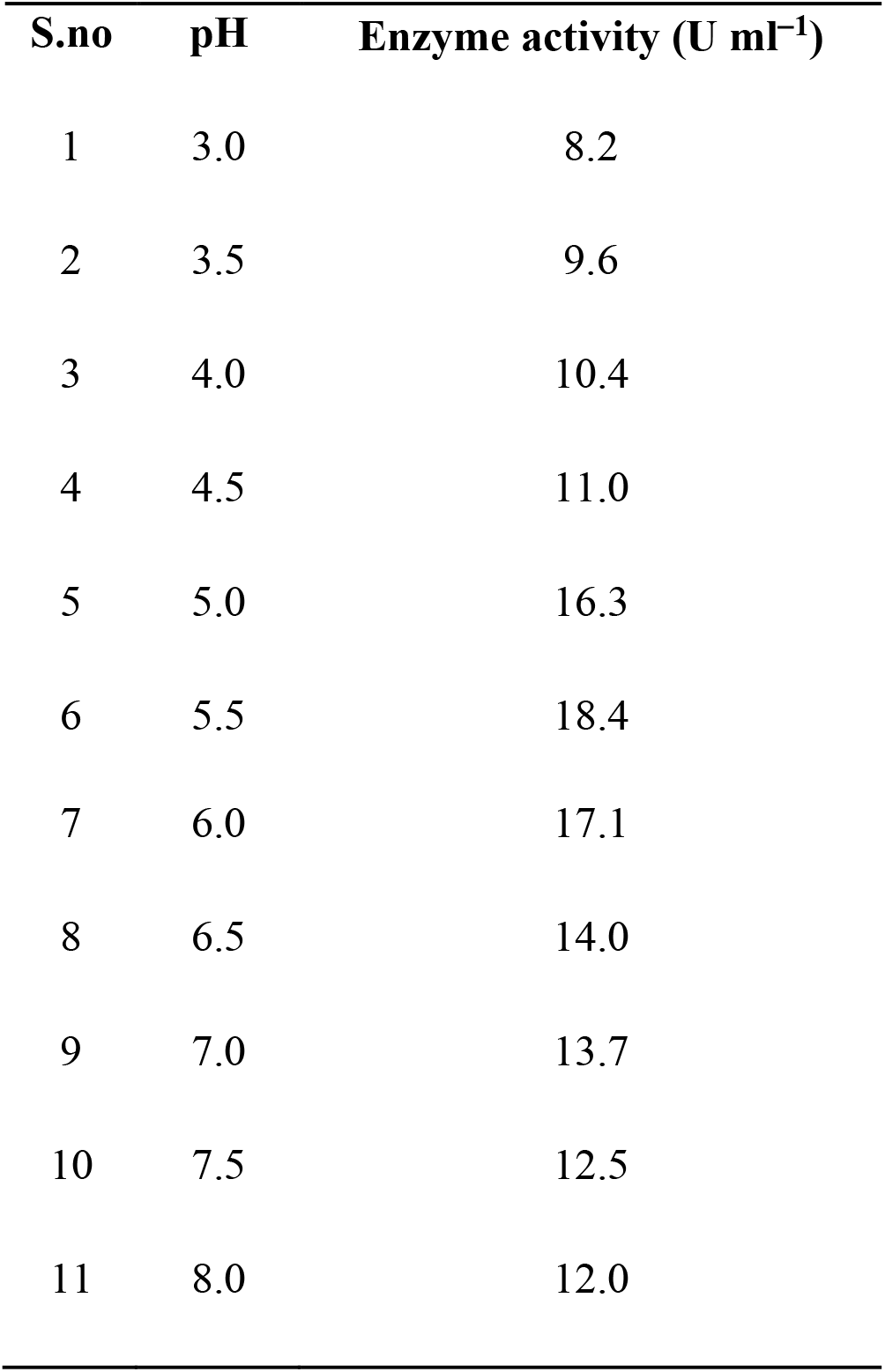
Effect of different pH on tannase activity

| S.no | pH | Enzyme activity (U ml<sup>-1</sup>) |
| --- | --- | --- |
| 1 | 3.0 | 8.2 |
| 2 | 3.5 | 9.6 |
| 3 | 4.0 | 10.4 |
| 4 | 4.5 | 11.0 |
| 5 | 5.0 | 16.3 |
| 6 | 5.5 | 18.4 |
| 7 | 6.0 | 17.1 |
| 8 | 6.5 | 14.0 |
| 9 | 7.0 | 13.7 |
| 10 | 7.5 | 12.5 |
| 11 | 8.0 | 12.0 |

#### Effect of temperature

Ellagic acid production was higher at temperature 30 °C which yielded tannases activity was 19.8 U ml^−1^ (Chart 4) and (Table 2). Previous study has stated that, optimum temperature for production of tannases at 30 °C using potent strains such as *Aspergillus oryzae*, *Penicillium chrysogenum* and *A. niger* [16]. Also studied the optimum temperature for production of tannases at 30 °C using *A. niger* [37]. Already reported that 12.1 mg/g of ellagic acid produced with the temperature 28 °C with *A. niger* from valonea tannins [28]. They have achieved 0.256 mg/l ellagic acid with the temperature 35 °C using *Aspergillus awamori* from mauha bark [31]. Contrarily, reported the 235.89 mg g^−1^ of ellagic acid from pomegranate at 60 °C. Some researchers reported that the rising of temperature would affect enzyme activity increased and denature the enzymes thus decrease activity [40].

**Table 2.** Effect of different temperature on tannase activity

| S.no | Temperature (°C) | Enzyme activity (U ml<sup>-1</sup>) |
| --- | --- | --- |
| 1 | 25 | 13.0 |
| 2 | 30 | 19.8 |
| 3 | 35 | 14.5 |
| 4 | 40 | 12.2 |
| 5 | 45 | 11.0 |
| 6 | 50 | 10.4 |
| 7 | 55 | 10.0 |

#### Effect of nitrogen source

Among various nitrogen sources used for EA production higher (25.6 U ml^−1^) with 0.2% sodium nitrate (NaNO_3_) than ammonium nitrate (21.2 U ml^−1^) and ammonium chloride (18.1 U ml^−1^). While other nitrogen sources such as sodium nitrite, potassium nitrate, ammonium sulfate and urea were found to affect the tannase production (Chart 5) and (Table 3). Highest tannase production was obtained with *P. variotii* was grown in ammonium nitrate as nitrogen source. Previous study observed the lower production of tannase with ammonium sulphate compared with ammonium nitrate could be due to the toxicity of sulphate ion itself on fungal growth [41].

**Table 3.** Effect of various nitrogen sources (0.2%) on tannase activity

| S.no. | Nitrogen sources (0.2%) | Enzyme activity (U ml<sup>-1</sup>) |
| --- | --- | --- |
| 1 | Sodium nitrate | 25.6 |
| 2 | Sodium nitrite | 13.2 |
| 3 | Potassium nitrate | 14.7 |
| 4 | Ammonium nitrate | 21.2 |
| 5 | Ammonium sulfate | 8.4 |
| 6 | Ammonium chloride | 18.1 |
| 7 | Urea | 9.3 |

#### Effect of carbon source

A concentration of 0.2% sugar from mango pulp favored both growth and enzyme production. Above 1% sugar provided, enzyme activity was decreased (Chart 6) and (Table 4). About 0.2% of glucose favoured both growth and enzyme production. However, above 1% glucose, growth and enzyme production were drastically affected due the osmotic stress of the medium [24]. One percent of carbon source was found to be optimum for tannase production [39].

**Table 4.** Effect of various concentration of sugar on tannase activity

| S. no. | Concentration of sugar (%) | Enzyme activity (U ml<sup>-1</sup>) |
| --- | --- | --- |
| 1 | 0.1 | 18.9 |
| 2 | 0.2 | 20.7 |
| 3 | 0.5 | 18.2 |
| 4 | 1.0 | 17 |
| 5 | 3.0 | 15.3 |
| 6 | 5.0 | 12.7 |

#### Production of ellagic acid and enzyme activity under optimized condition

The ellagic acid production was carried out under optimized conditions (pH, temperature, carbon, nitrogen) and using mango pulp waste. The enzyme activity was calculated to be as follows (Table 5). Already reported the enzyme activity of *Aspergillus oryzae* was co-cultured with *Trichoderma reesei* using acorn cups extract containing up to 62% ellagitannins as substrate to produce ellagic acid with relatively high levels of ellagitannin acyl hydrolase, cellulase and xylanase [42]. The results indicate that the mixed culture is an effective approach to produce an enzyme system of degrading ellagitannins for ellagic acid production.

**Table 5.**
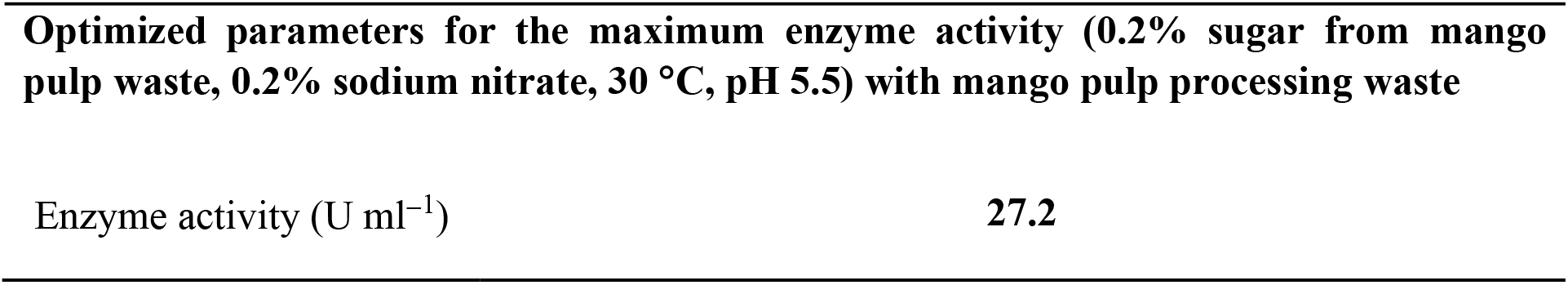
Optimized parameters for Enzyme activity

## Conclusion

Ellagic acid produced through bioconversion has gained great attention in recent years due to their economic and ecofriendly nature. EA production from the mango pulp waste extract using *A. niger* had produced a 200 μg of EA from 3% g of mango pulp waste at the optimum conditions (0.2% sugar, 0.2% sodium nitrate, 30 °C and pH 5.5). The present study was considered as a pioneer attempt to synthesis the ellagic acid from the mango pulp waste using GRAS stain. It is promising that smaller industries can improve their productivity with bioconversion of the waste generated in the process. Solid state production of ellagic acid using *Aspergillus niger* would be cost-effective and safe method producing pharmaceutically important ellagic acid.

**Chart 1:**
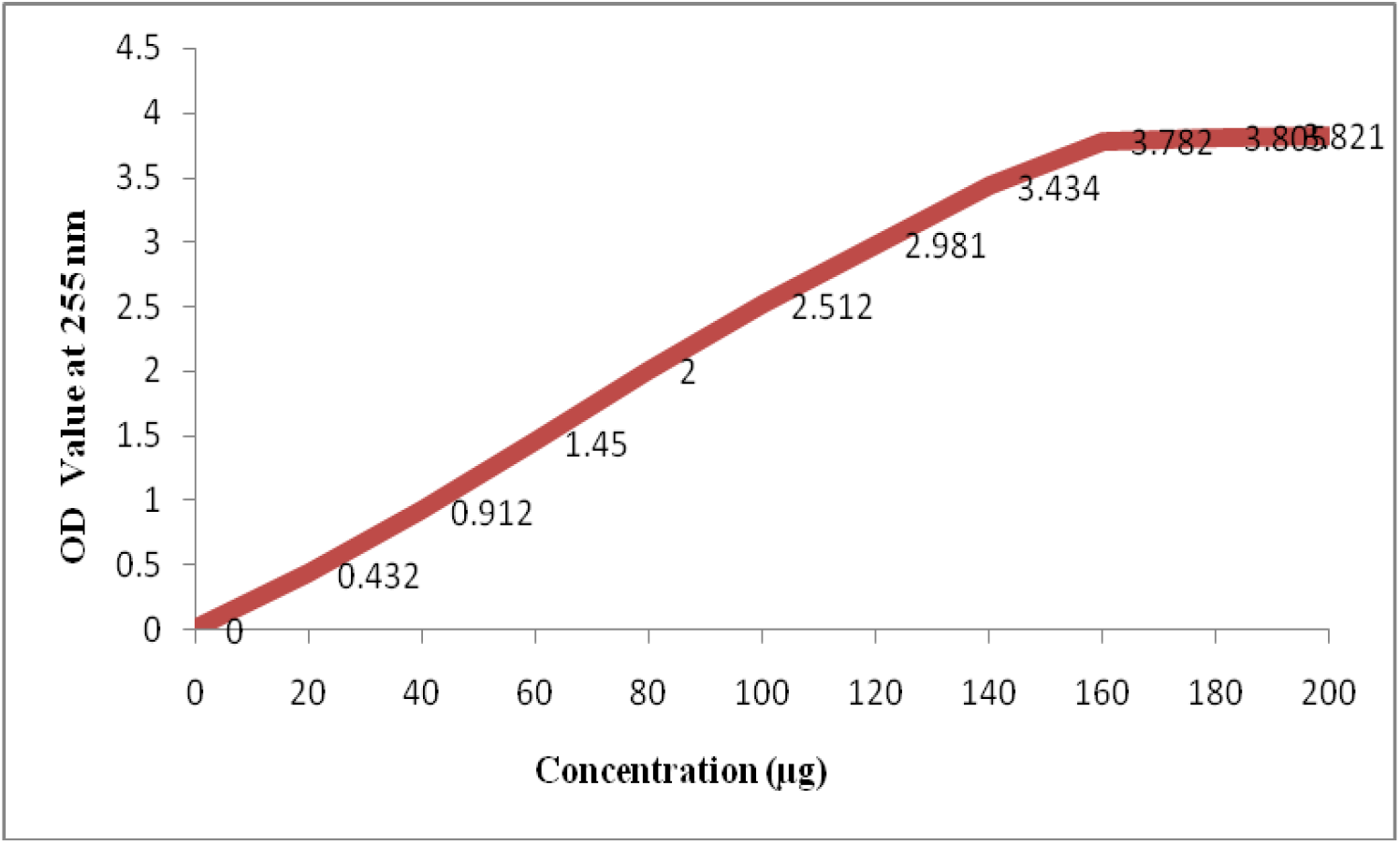
Quantification of ellagic acid

**Chart 2:**
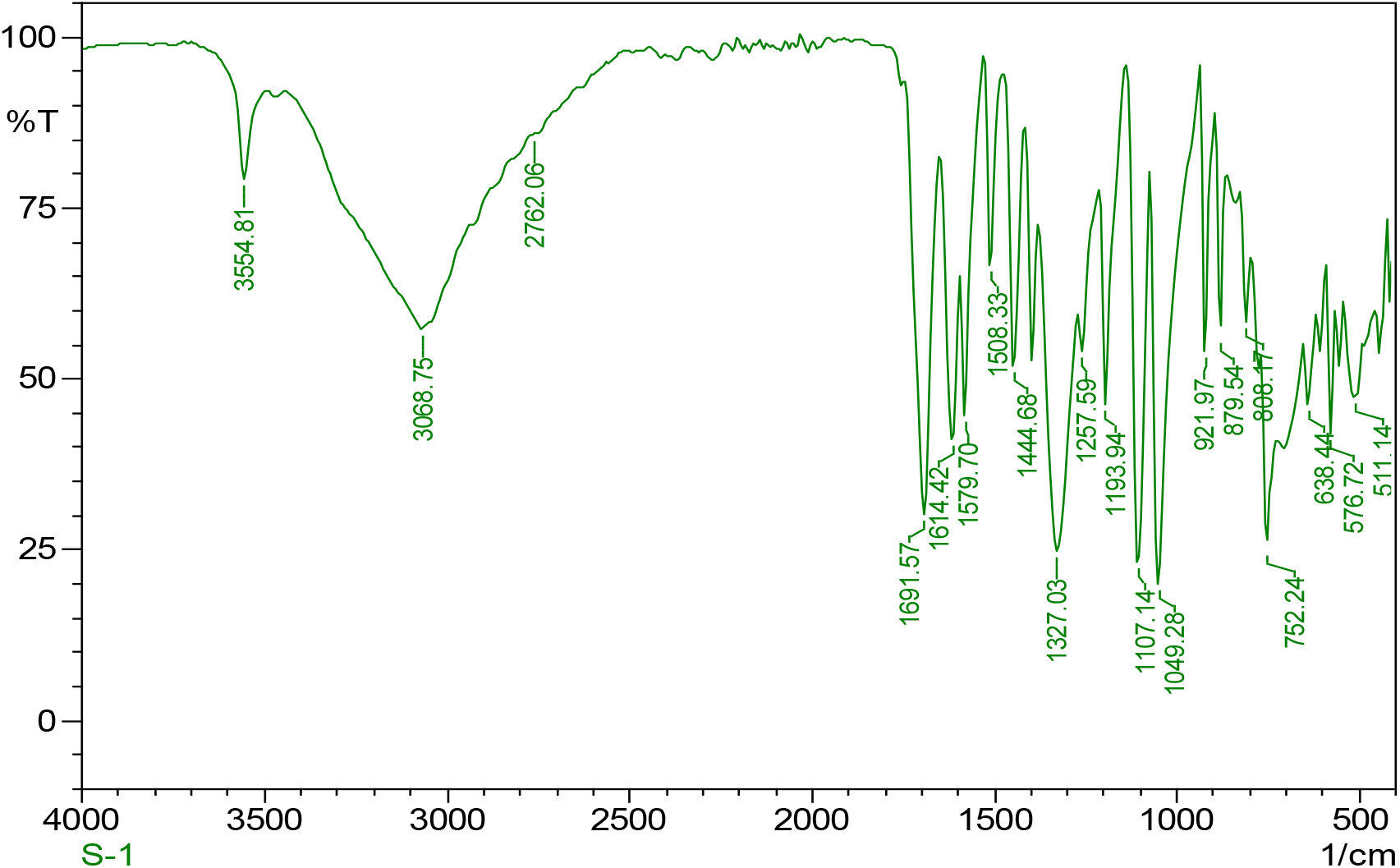
FTIR spectrum of sample

**Chart 3:**
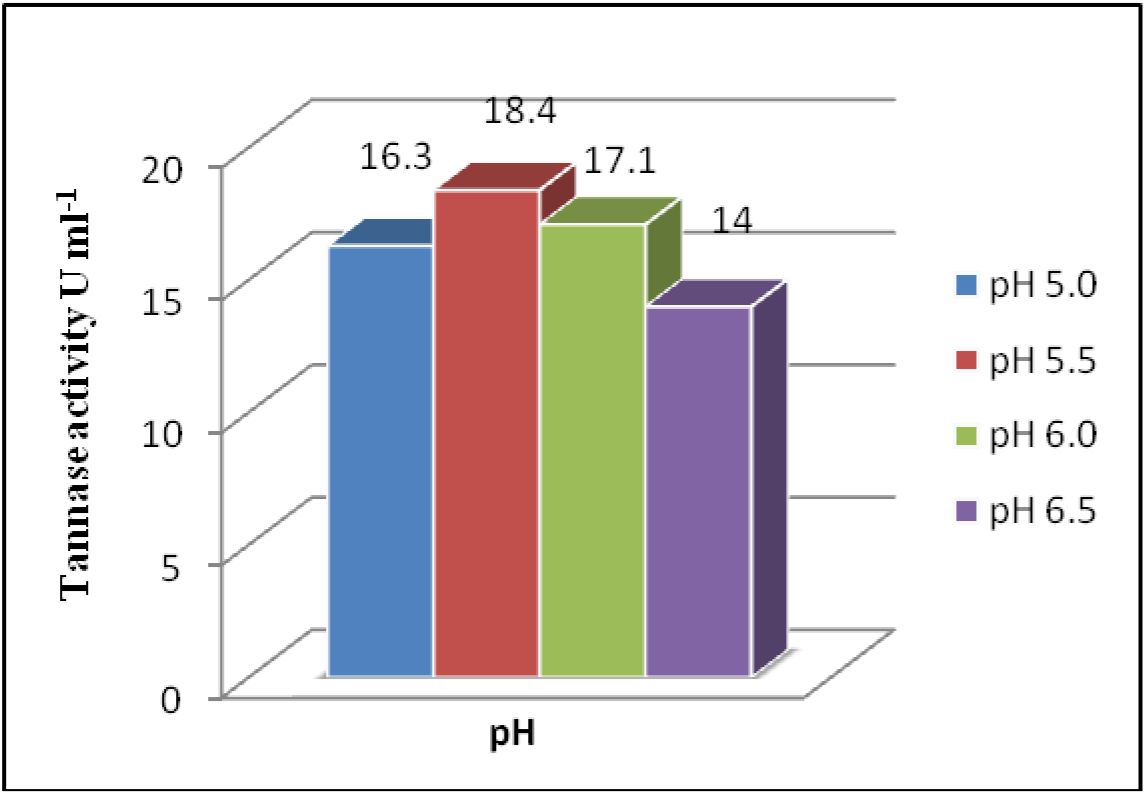
Effect of pH on tannase activity

**Chart 4:**
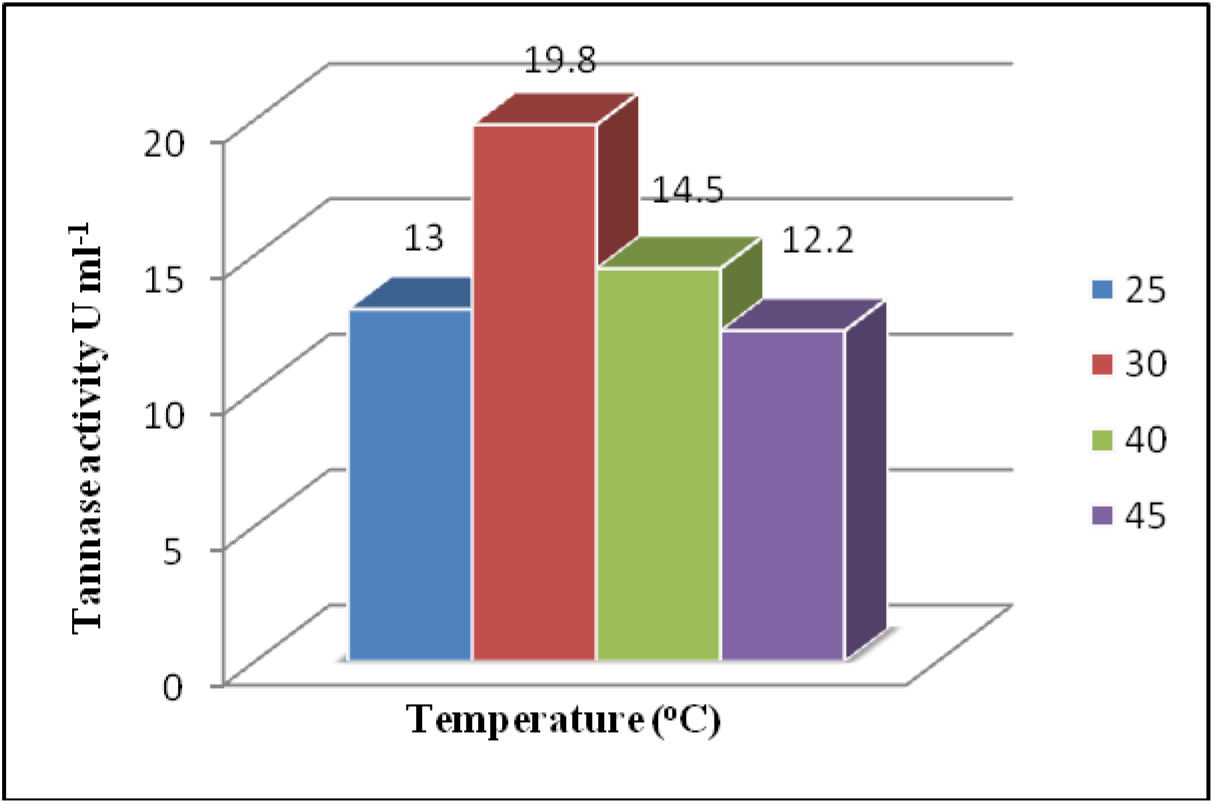
Effect of temperature on tannase activity

**Chart 5:**
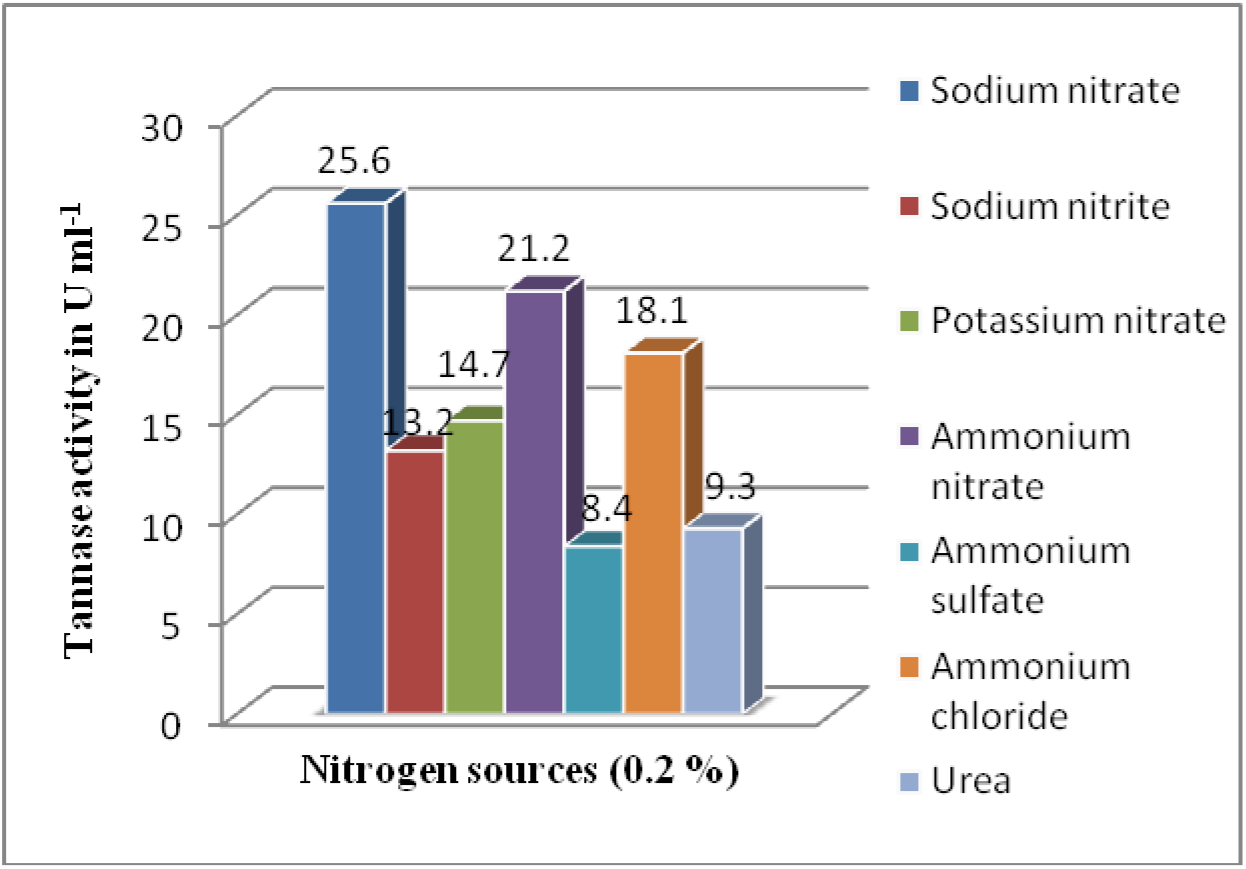
Effect of nitrogen sources on tannase activity

**Fig 6:**
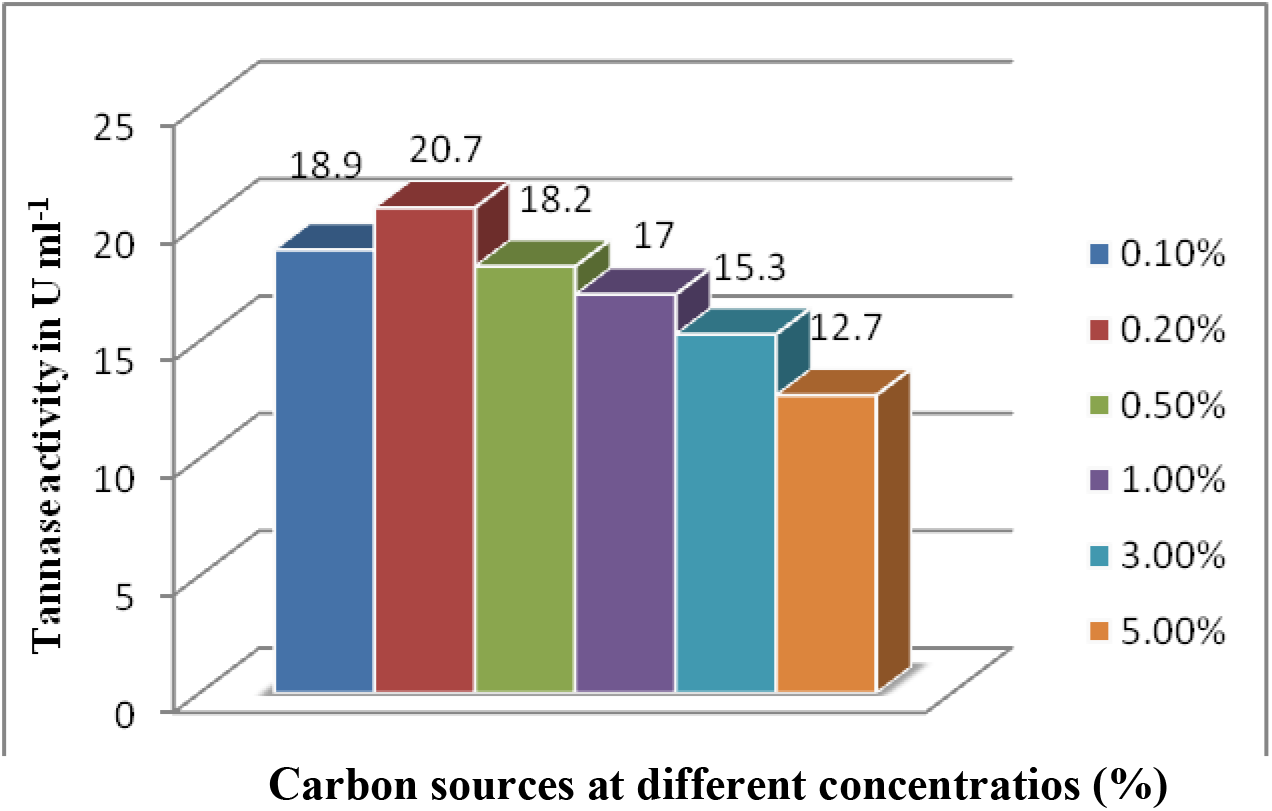
Effect of different concentration of carbon sources on tannase activity

## Abbreviations used

EA: Ellagic acid
*A.niger*: *Aspergillus niger*
GRAS: generally recognized as safe
CFR: Code of federal regulations
PSH: Pomegranate seeds and husk
PDA: Potato Dextrose Agar
LPCB: Lactophenol Cotton Blue
OD: optical density
EI: electron impact
SSF: solid state fermentation
FTIR: Fourier-transform infrared spectroscopy

## Acknowledgment

Both the authors acknowledge the DST-FIST facilities (Ref No. SR / FST / LSI – 640 / 2015C).

## Notes

The authors declare no competing financial interest.

